# Acculturation stressors and academic adjustment among Nepalese students in South Korean higher education institutions

**DOI:** 10.1101/2020.12.07.414441

**Authors:** Madhu S. Atteraya

## Abstract

**Introduction:** International students are steadily increasing in South Korean higher education institutions. How well international students in South Korea are adjusted academically and the relationship between acculturation stressors and academic adjustment has not yet been determined. The study aimed to fill this research gap by examining the relationship between acculturation stressors and academic adjustment among Nepalese international students in South Korean higher education institutions.

**Methods:** The sample of the study consisted of Nepalese international students who enrolled in 36 universities in South Korea. Students’ background characteristics and acculturation stressors (e.g., discrimination, homesickness, hate/rejection, fear, cultural shock, and guilt) were selected to assess the association of these characteristics and stressors with academic adjustment. Pearson correlation and hierarchical multiple regression were utilized to examine the association between acculturation stressors and academic adjustment.

**Results:** The results from the Pearson correlation revealed the negative correlation of perceived discrimination (r = -.23, p< 0.01), perceived hate/rejection (r = -.18, p< 0.05), perceived fear (r = -.24, p< 0.01), and perceived cultural shock (r = -.17, p< 0.05) with academic adjustment. Further, the hierarchical regression model revealed that marital status (*β* = .223, *p <*.01) had a positive association with academic adjustment, whereas perceived fear (*β* = -.206, *p <*.05) had a negative association with academic adjustment even after including students’ background characteristics and other acculturation stressors.

**Conclusion:** Addressing acculturation stressors among international students in higher education institutions is essential to facilitate positive academic adjustment. Mainly, perceived fear has negatively affected students’ academic adjustment. Based on these findings, tailored programs must be developed to curtail students’ perceived fear in order to enhance their academic performance in South Korean higher education institutions.

## Introduction

International students’ social and ethnic identities, backgrounds, communication skills, and acculturation levels and the effects of these aspects on their academic performance and adjustment in host cultures have become an essential concern in higher education institutions worldwide [1]. Evidence from Western higher education institutions has suggested that international students experience a higher level of psychological, social, and academic distress [2-5]. Therefore, the extent to which acculturation stressors affect academic adjustment among international students in different cultural settings (i.e., non-Western contexts) requires similar scrutiny in order to better adapt the increasing number of international students to their host culture, such as the Republic of Korea (henceforth South Korea).

Acculturation is a critical factor that affects immigrants’ health status (including psychological, somatic, and social aspects) while they are in the process of adjusting to a dominant culture [6]. Furthermore, acculturation stressors cause a high level of distress for an individual in achieving desired outcomes in new cultural settings [6]. For international students from non-English-speaking countries with lower socioeconomic status, they have increased difficulty in academic adjustment due to language-related barriers, cultural differences, and economic hardships. These students experience a significant level of acculturation stress, suffer from poor physical and mental health, and encounter challenges in academic achievement [7–9]. For instance, in the United States of America, a growing number of studies demonstrated that ethnic minority students have a higher level of acculturation stress and difficulty in academic adjustment to mainstream US educational institutions. Evidence suggested that Latina/o and Native American students are disadvantaged groups with lower academic adjustment potential due to their cultural heritage, speaking English in their native accent or dialects, and their sense of exclusion from US educational institutions [10–13]. Similar findings in Australia and Europe report that a substantial number of students expressed distress, felt unconnected, encountered lower levels of social and academic integration, and experienced poor emotional wellbeing [4].

Several aspects have been revealed as risk factors for academic adjustment among international students. Studies have found these risk factors to include cultural norms, language barriers, perceived discrimination, financial problems, work restrictions, higher tuition fees, accommodation, and transportation problems [5]. For example, Turkish Muslim students who participated in the Erasmus student exchange to a European country had difficulty adjusting due to fear of the unknown, loneliness, food shock, and being an object of suspicion [2]. These risk factors, or stressors, further exacerbate international students’ acculturation stress, impacting their academic adjustment capability and performance in higher education institutions.

Abundant evidence from previous studies has shown that among immigrants, refugees, and international students, there is an association of acculturation stressors with ill (mental) health status, substance dependency, social maladaptation, and hindrances to achieving desired preference in a host society[14]. For instance, a study conducted among Korean international students living in the Pittsburgh area found that acculturation stressors were strongly correlated to poor mental health status [15].

### Context of South Korea

With a population of around 51.63 million, South Korea historically emphasized educational attainment for its citizens even in times of economic hardship. By the end of the Korean War (1950–1953), the Korean Peninsula gradually overcame its absolute poverty and has now transformed into a prosperous country. Today, South Korea is known as the “Asian Tiger,” ranking third in Asia and 13^th^ in the world economy. The socioeconomic transformation from poverty-stricken society to prosperity is a result of Confucian values of diligence and importance given to educational attainment and rapid expansion of higher education institutions [16].

Economic advancement in South Korea led to the internationalization of South Korean higher education institutions, adding to Korean higher education’s global competency. Consequently, South Korea experienced a quantitative expansion of international students, as there were 4,000 international students in 2004, which increased to 85,923 in 2011 [17]. As of January 2020, a total of 118,342 international students were in South Korea, the majority of them from Asia, including China (59,720), Vietnam (18,640), Mongolia (5,788), Nepal (1,964), and Japan (1,919) – ranking Nepal as the fourth-largest contributor to the population of international students in South Korea [18]. Moreover, the South Korean government has announced a plan to increase the number of international students to 200,000 by 2023 with the goal of making South Korea an educational hub in Asia.

Along with the increasing number of international students in South Korea and positioning South Korea as an educational hub, the South Korean government attempted to improve academic competencies via various policy innovations, including Brain Korea 21 Project, World Class University Project, Humanity Korea, Social Science Korea, University for Creative Korea, Brain Korea 21, and BrainKorea21 Plus [19]. Additionally, the National Research Foundation of Korea provides various research funds to assist university professors and students (including international students) in enhancing their academic performance and competency in South Korean higher education institutions. Specifically, South Korea attempts to extend itself in knowledge economics by strengthening its academic competitiveness at different levels (including at the university, faculty, and student level). To do so, the country invested US$73.3 billion for research and development to strengthen higher education institutions to a “global standard” [20]. Along with all these efforts, how well international students are adjusted in South Korean academic institutions is an essential concern for policymakers and educational researchers.

Jon [21] recognized that with the increasing number of international students in South Korea, international students’ academic adjustment is critical to ensuring that Korean students may develop intercultural learning. However, international students’ academic adjustment issues in South Korea have not been well documented. Specifically, most relevant studies were conducted in the Korean language and emphasized Chinese students’ acculturation issues; of these studies, results were mixed in terms of acculturation levels [22-24]. For example, Lee, Jon, and Byun [24] mentioned that Chinese students in South Korea were less accepted, felt discriminated against, and experienced negative stereotypes in comparison to students from North America and Europe. In contrast, Jon, Lee, and Byun [23] revealed that Chinese students think of South Korea as an attractive destination due to scholarship opportunities, geographical proximity with China, employability after graduation, safety and security, and easier visa accessibility than Japan and other Western countries. Consequently, Chinese students expressed increased academic satisfaction in terms of academic resources, facilities, and quality of instruction in South Korean higher education institutions. Similarly, Alemu and Cordier [25] demonstrated that international students are generally satisfied in South Korean higher education institutions.

However, South Asian international students’ academic adjustment and integration in South Korea has not yet been considered outside of a study by Bhandari [26], which analyzed the association between acculturation stressors and health-related quality of life. The current study attempts to fill this research gap by examining the acculturation stressors experienced by South Asian international students’ in South Korea. More specifically, the study examines the extent to which certain acculturation stressors affect Nepalese international students’ academic adjustment in South Korean higher education institutions.

## Methods

### Procedure

The researcher distributed questionnaires to Nepalese students during the 12^th^ Annual General Meeting (AGM) of a Nepalese students’ organization called the Society of Nepalese Students in South Korea (SONSIK). The SONSIK’s 12^th^ AGM was organized at Sunmoon University, Asan, Korea, on 22^nd^–23^rd^ of August 2015. The questionnaires were distributed before the formal program started on the 22^nd^ of August, and the researcher collected them on another day after the end of the program. All the questionnaires were administered in the English language.

### Participants

Most Nepalese students attended the SONSIK AGM, as the SONSIK executive committee members asked all Nepalese students to attend the annual meeting. The AGM is an opportunity to meet friends at one place once in a year. Of these Nepalese students, 174 from 36 universities in South Korea participated in the study. Among them, 136 students returned the self-administered questionnaire. Four cases were removed due to missing information. Ultimately, a sample size of 132 was analyzed. In 2015, a total of 580 Nepalese students enrolled in all South Korean universities [27]. Therefore, the study sample is acceptable for data analysis with 7.36% margin of error at the 95% confidence interval.

### Ethics Statement

Ethical consideration was concerned for the data collection process to maintain the respondents’ privacy, confidentiality, and voluntary participation. For instance, in the cover letter of the questionnaire, the researcher mentioned the following statement — “Your information will remain confidential; please do not include your name. If you choose to participate, please answer all questions as honestly as possible and return the completed questionnaire promptly.

Participation is strictly voluntary.” The questionnaires were placed at a specific place to maintain privacy, confidentiality, and voluntary participation and asked to return the questionnaires voluntarily another day. Adhering to these ethical considerations, questionnaires were collected on another day. The questionnaires were anonymized, and the data collection process was completely voluntary.

### Instruments

#### Acculturative stressor

The researcher used the Acculturative Stress Scale for International Students (ASSIS), created by Sandhu and Asrabadi [28]. The ASSIS consists of 36 items in total, including perceived discrimination (13 items), homesickness (4 items), perceived hate/rejection (5 items), fear (4 items) and stress due to cultural change (3 items), guilt (2 items), and non-specific concerns (5 items). A 5-point Likert scale ranging from 1 (strongly disagree) to 5 (strongly agree), where a higher score represents a higher level of acculturative stress, was incorporated. The reliability Cronbach’s alpha for the 36-item scale was .92; the reliability Cronbach’s alpha for each stressor ranged from .70 to .92.

#### Academic adjustment

The researcher used Baker and Siryk’s [29] scale to measure academic adjustment with slight modifications. The researcher created Cluster 2 questions that referred to students’ motivation, while Cluster 3 referred to students’ performance. Then, Cluster 2 and Cluster 3 were combined to measure students’ academic adjustment. Cluster 2 had the questions as: I am definite about reasons for being in Korea to study; I have well-defined academic goals; I consider a college/university degree important; I enjoy academic work, my interests are related to current research work; I doubt the value of college/university degree in Korea. Cluster 3 included: I find academic work difficult; I do not function well during exams or experiments; I am not satisfied with academic performance; I do not feel smart enough for course/research work; I do not use study time efficiently; I do not enjoy writing papers for courses/research; I have trouble concentrating when studying; I do not do well academically considering the effort; I have trouble getting started on homework. The rating scale was Yes, No, and Do Not Know. Positive attitudes on academic adjustment were coded as 1, and non-positive attitudes towards their academic adjustment were coded as 0. The reliability Cronbach’s alpha for the academic adjustment scale was .75.

### Data Analysis

First, descriptive statistics were performed. Then, Pearson correlation testing was conducted to examine the associations between students’ background characteristics, the five dimensions of acculturative stress (e.g., discrimination, homesickness, hate/rejection, fear, cultural shock, and guilt), and students’ academic adjustment. Finally, hierarchical multiple regression was conducted using SPSS.

## Results

Table 1 presents the demographic information of Nepalese students’ who were enrolled in 36 Korean higher education institutions. Most students (78.8%) were male, whereas only 21.2 % of students were female. There was an almost equal proportion of married (48.5%) to never married (51.5%) students. Approximately 53.8% of students did not feel comfortable using the Korean language in everyday communication. Most of the students (72.7%) received a full scholarship to pursue their higher education in South Korea.

**Table 1.**
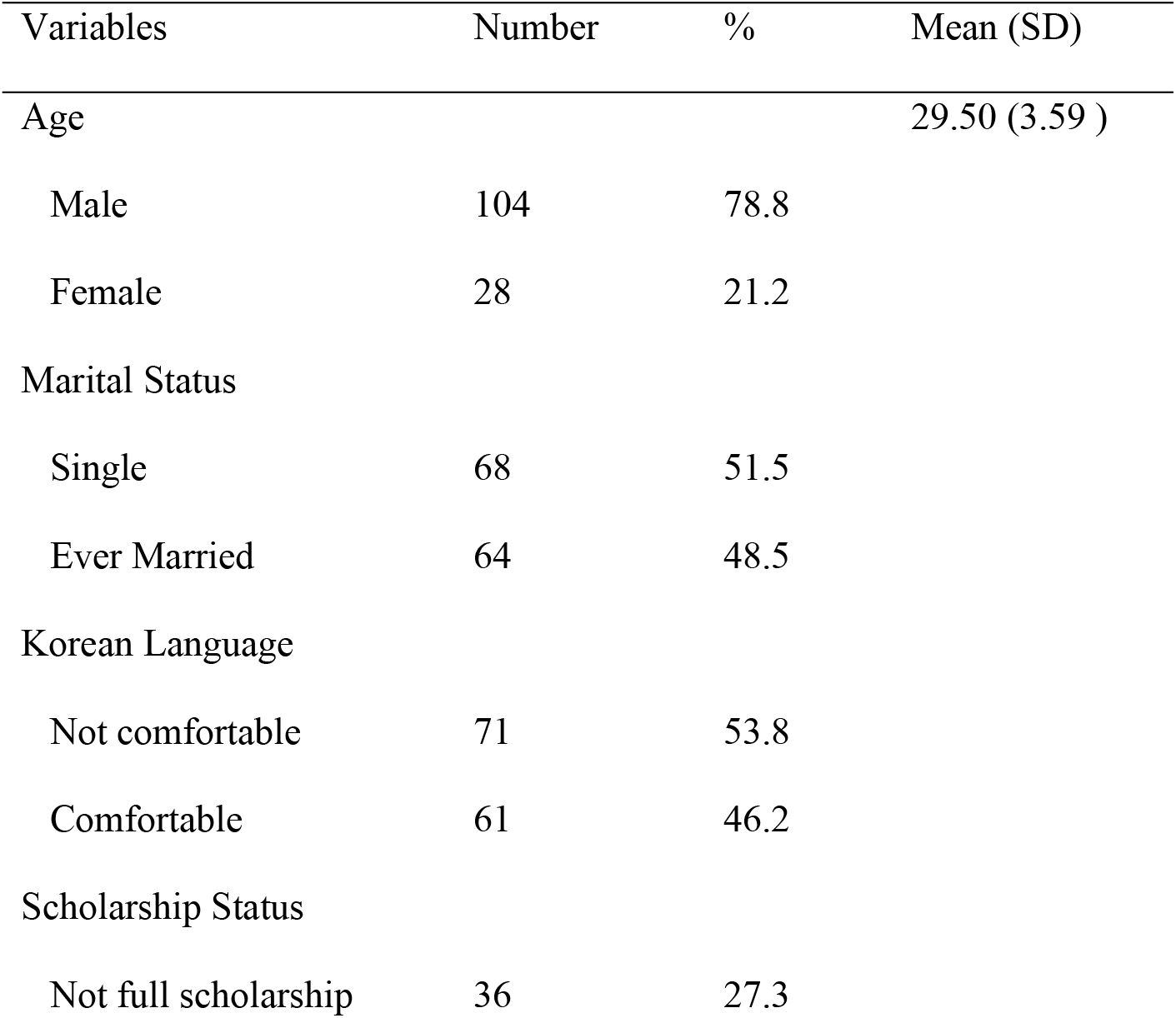
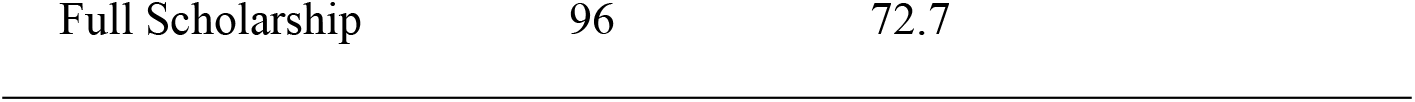
Demographic Information of the Sample (Sample = 132)

| Variables | Number | % | Mean (SD) |
| --- | --- | --- | --- |
| Age |  |  | 29.50 (3.59 ) |
| Male | 104 | 78.8 |  |
| Female | 28 | 21.2 |  |
| Marital Status |  |  |  |
| Single | 68 | 51.5 |  |
| Ever Married | 64 | 48.5 |  |
| Korean Language |  |  |  |
| Not comfortable | 71 | 53.8 |  |
| Comfortable | 61 | 46.2 |  |
| Scholarship Status |  |  |  |
| Not full scholarship | 36 | 27.3 |  |

Table 2 presents the correlation for background characteristics, acculturation stressors, and academic adjustment. As shown in Table 2, students’ full scholarship (r = -.18, p< 0.05) was negatively correlated with the level of Korean language comfort level. Homesickness (r = .17, p< 0.05) had negative correlation with Korean language comfort level. Homesickness (r = .22, p< 0.01) and perceived discrimination had positive correlation. Similarly, perceived hate/rejection (r = .75, p< 0.01), fear (r = .38, p< 0.01), cultural shock (r = .31, p< 0.01), and guilt (r = .20, p< 0.05) had positive correlation with perceived discrimination, respectively. Perceived hate/rejection (r = .30, p< 0.01), cultural shock (r = .36, p< 0.01, and guilt (r = .37, p< 0.01) had positive correlation with homesickness, respectively. Perceived fear, cultural shock, and guilt had positive correlation. Moreover, academic adjustment (r = .19, p< 0.05) had positive association with married students. However, academic adjustment (r = -.23, p< 0.01) had negative correlation with perceived discrimination (r = -.23, p< 0.01). Academic adjustment (r = -.18, p< 0.05) had negative correlation with perceived hate/rejection. Academic adjustment (r = - .24, p< 0.01) had negative correlation with perceived fear. Similarly, academic adjustment (r = - .17, p< 0.05) had negative correlation with cultural shock.

**Table 2.** Correlation of Statistical Control, Independent, and Dependent Variables

| Variables | 1 | 2 | 3 | 4 | 5 | 6 | 7 | 8 | 9 | 10 | 11 |
| --- | --- | --- | --- | --- | --- | --- | --- | --- | --- | --- | --- |
| 1. Female | — |  |  |  |  |  |  |  |  |  |  |
| 2. Married | 0.12 | — |  |  |  |  |  |  |  |  |  |
| 3. Korean language | 0.03 | -0.04 | — |  |  |  |  |  |  |  |  |
| 4. Full scholarship | -0.14 | -0.08 | -.18* | — |  |  |  |  |  |  |  |
| 5. Discrimination | -0.07 | 0.03 | -0.13 | 0.05 | — |  |  |  |  |  |  |
| 6. Homesickness | 0.09 | 0.03 | -.17* | 0.07 | .22** | — |  |  |  |  |  |
| 7. Hate/Rejection | -0.08 | 0.04 | -0.12 | 0.07 | .75** | .30** | — |  |  |  |  |
| 8. Fear | -0.09 | 0.06 | -0.08 | -0.01 | .38** | 0.14 | .48** | — |  |  |  |
| 9. Culture shock | 0.01 | -0.06 | -0.12 | 0.13 | .31** | .36** | .26** | .20* | — |  |  |
| 10. Guilt | -0.15 | 0.05 | -0.05 | -0.05 | .20* | .37** | .32** | .23** | .28** | — |  |
| 11. Academic Adjustment | -0.01 | .19* | -0.02 | -0.00 | -.23** | -0.16 | -.18* | -.24** | -.17* | -0.13 | — |

Note: \* $p < 0.05$ . \*\* $p < 0.01$

Table 3 presents the hierarchical linear regression model with students’ background characteristics and acculturation stressors predicting academic adjustment. Model 1 (R2 = .041) presents the effect of students’ background characteristics (e.g., gender, marital status, Korean language use, and full scholarships) on academic adjustment. The results revealed that marital status (*β* = .202, *p <*.05) was positively associated with academic adjustment. However, other background characteristics (i.e., gender, Korean language use, and full scholarships) were not associated with academic adjustment. Model 2 included all background characteristics and academic stresses to examine their effects on academic adjustment. As such, Model 2 (R2 = .157) demonstrates that marital status (*β* = .203, *p <*.01) remained a positive association with academic adjustment. However, perceived fear (*β* = -.206, *p <*.05) was negatively associated with academic adjustment. Other acculturation stressors (i.e., discrimination, homesickness, hate/rejection, cultural shock) did not have a statistically significant effect on academic adjustment.

**Table 3.** Hierarchical Linear Regression Analyses

| Variable | Model 1 |  |  | Model 2 |  |  |
| --- | --- | --- | --- | --- | --- | --- |
| | *B* | *SE B* | $\beta$ | *B* | *SE B* | $\beta$ |
| Constant | 9.438 | .740 |  | 13.859 | 1.397 |  |
| Gender | -.267 | .695 | -.034 | -.456 | .694 | -.058 |
| Marital Status | 1.298 | .566 | .202* | 1.430 | .549 | .223** |
| Korean language | -.081 | .571 | -.013 | -.456 | .559 | -.071 |
| Full scholarship | .016 | .646 | .002 | .014 | .633 | .002 |
| Discrimination |  |  |  | -.058 | .035 | -.215 |
| Homesickness |  |  |  | -.085 | .083 | -.101 |
| Hate/Rejection |  |  |  | .073 | .088 | .115 |
| Fear |  |  |  | -.178 | .084 | -.206* |
| Culture shock |  |  |  | -.042 | .095 | -.042 |
| Guilt |  |  |  | -.076 | .134 | -.055 |
| *R*<sup>2</sup> |  | .041 |  |  | .157 |  |
| *Adjusted R*<sup>2</sup> |  | .010 |  |  | .087 |  |
| *R*<sup>2</sup> change |  | .041 |  |  | .116 |  |
| *F* for change in *R*<sup>2</sup> |  | 1.343 |  |  | 2.774* |  |

Note: \**p* < .05. \*\**p* < .01.

## Discussion

Understanding the factors that affect academic adjustment among international students is essential for enhancing academic competitiveness. In order to strengthen international students’ academic adjustment capabilities and the academic competency of higher education institutions, policymakers and practitioners must be well-informed of these factors. However, to the best of the researcher’s knowledge, studies on these factors as barriers toward academic adjustment have not yet been conducted in a South Korean context.

Academic adjustment is crucial, as students with better academic adjustment capabilities possess a higher level of personal satisfaction and academic success, yielding better output in a competitive educational environment. In South Korea, universities are experiencing increasing numbers of international students, with most universities in South Korea gradually heading toward global competitiveness, thereby enhancing South Korea universities’ global ranking. Moreover, at the macro level, internationalization of higher education in South Korea functions to strengthen “soft power,” enriching Korean culture and the academic system globally. In such a context, it is essential to understand the extent to which acculturation stressors impede or facilitate academic adjustment among international students in the South Korean educational environment. Here, the key findings of the study have elucidated Nepalese international students’ academic adjustment in South Korean higher education institutions

First, the study revealed that marital status had a positive association for academic adjustment among Nepalese students in South Korean higher education institutions. The result is inconsistent with previous studies in the Asian context, which, in the case of Malaysian higher education institutions, found that international students’ marital status did not influence sociocultural adjustment [30]. Similar findings have revealed that international academics’ marital status did not influence sociocultural adjustment in Saudi Arabian higher education institutions either [31]. Additionally, inconsistent with other studies’ findings [13,32], married Asian students have a lower academic adjustment level in the USA and German universities. However, in the current study, both in bivariate and multivariate hierarchical regression analyses, students’ marital status remained a robust positive predictor for academic adjustment. This may suggest that married Nepalese students in South Korea are more motivated and determined to work toward academic achievement.

Second, in the bivariate level, the study revealed that perceived discrimination, hate/rejection, fear, and cultural shock had negatively correlated with academic adjustment among Nepalese students in South Korean higher education institutions. Specifically, perceived discrimination (r = -.23) and fear (r = -.24) had strong negative correlation (**p< 0.01) with academic adjustment. Even in the USA, international students experience various discrimination levels based on the regions (or nations) they come to the USA [33]. The level of perceived discrimination negatively impacted educational experience and actual academic performance [34]. Likewise, Nepalese students may perceive themselves as being discriminated against in the South Korean cultural context. Consequently, those who perceived themselves as being discriminated against had their self-esteem negatively affected, thereby leading to decreased competence for educational attainment, including academic adjustment. Indeed, perceived discrimination leads to social and academic marginalization from the mainstream academic culture, such as deterring student engagement from learning outcomes.

Third, perceived fear remained the strongest predictor that negatively affected Nepalese international students’ academic adjustment in South Korea in bivariate and hierarchical linear regression analysis. Although a further qualitative study is required to determine the factors that trigger “fear” among international students in South Korea, the study confirmed that perceived fear ultimately negatively affected Nepalese students’ academic adjustment. The findings may imply several reasons for triggering this fear, such as the fear of uncertainty in new Korean educational and sociocultural contexts, language changes and cultural differences hindering effective communication, fear of failure, fear of making mistakes, fear of non-native accents, and the fear for the unknown. Perhaps in academia, the hierarchical social structure makes it more challenging to adjust, and that too may create “fear” for international students.

Fourth, the study revealed that perceived fear remained a strong predictor for academic adjustment negatively in the hierarchical linear model. Similarly, perceived discrimination, hate/rejection, fear, and cultural shock had negatively affected the academic adjustment at the bivariate level. Higher education institutions have an awareness of students’ perceived difficulties, such that universities have initiated several programs to support students’ mental health issues. For example, currently, several universities have support centers for international students’ mental health through university-centered counseling programs concentrating on topics related to depression, anxiety, emotional difficulties, academic stress, and personal and relationship problems. The findings of the study further provide insight to counseling centers that they must help international students in overcoming perceived fear. However, most international students are unlikely to use university-centered counseling services due to language and cultural barriers since most services are provided in the Korean language. There are some testimonies of international students who expressed their difficulties effectively communicating with counselors who deliver their services in the Korean language.

Fifth, the current study has some limitations. The study used cross-sectional data and only employed Nepalese international students in the South Korean cultural context. Further study is needed to explore adaptation strategies and difficulties in Korean academic culture [35]. An additional qualitative study needs to dig deeper into reasons for exploring academic adjustment barriers among students from diverse sociocultural backgrounds, as there are several international students from various nations in any given university. Another limitation of the study is that the study is limited to Nepalese international students’ experiences. Despite this constraint, the study’s findings have merit as they identified acculturation stressors (i.e., perceived fear) as crucial barriers to academic adjustment. This result helps inform national policies to better adapt international students to South Korean higher education with ease and grace. Well-adapted students and students with fewer acculturation problems could contribute more effective and successful results in a highly competitive educational environment.

In conclusion, addressing acculturation stressors among international students may yield higher levels of personal satisfaction, enhance productive academic life, and increase performance among international students living and studying in South Korea. Doing so ultimately enables students to strengthen Korean higher education institutions’ academic competitiveness in the long run.

## References

1. Hawkes L. The development of the social and academic identities of international students in English-speaking higher education institutions. York St John University School of Foundation and English Language Studies BPP University. 2014.

2. Brown L, Aktas G. Fear of the unknown: a pre-departure qualitative study of Turkish international students. British Journal of Guidance & Counselling. 2011;39(4):339–55.

3. Fritz MV, Chin D, DeMarinis V. Stressors, anxiety, acculturation and adjustment among international and North American students. International Journal of Intercultural Relations. 2008;32(3):244–59.

4. Rienties B, Beausaert S, Grohnert T, Niemantsverdriet S, Kommers P. Understanding academic performance of international students: The role of ethnicity, academic and social integration. Higher education. 2012;63(6):685–700.

5. Smith RA, Khawaja NG. A review of the acculturation experiences of international students. International Journal of intercultural relations. 2011;35(6):699–713.

6. Berry JW, Kim U, Minde T, Mok D. Comparative studies of acculturative stress. International migration review. 1987;21(3):491–511.

7. Hamamura T, Laird PG. The effect of perfectionism and acculturative stress on levels of depression experienced by East Asian international students. Journal of Multicultural Counseling and Development. 2014;42(4):205–17.

8. Mori SC. Addressing the mental health concerns of international students. Journal of Counseling & Development. 2000;78(2):137–44.

9. Poyrazli S, Kavanaugh PR, Baker A, Al-Timimi N. Social support and demographic correlates of acculturative stress in international students. Journal of College Counseling. 2004;7(1):73–82.

10. Acevedo-Polakovich ID, Quirk KM, Cousineau JR, Saxena SR, Gerhart JI. Acting bicultural versus feeling bicultural: Cultural adaptation and school-related attitudes among US Latina/o youth. Journal of Hispanic Higher Education. 2014;13(1):32–47.

11. Fischer S, Stoddard C. The academic achievement of American Indians. Economics of Education Review. 2013;36:135–52.

12. Fryberg SA, Covarrubias R, Burack JA. Cultural models of education and academic performance for Native American and European American students. School Psychology International. 2013;34(4):439–52.

13. Souto-Manning M. Competence as linguistic alignment: Linguistic diversities, affinity groups, and the politics of educational success. Linguistics and Education. 2013;24(3):305–15

14. Nguyen A-MD, Benet-Martínez V. Biculturalism and adjustment: A meta-analysis. Journal of Cross-Cultural Psychology. 2013;44(1):122–59.

15. Lee J-S, Koeske GF, Sales E. Social support buffering of acculturative stress: A study of mental health symptoms among Korean international students. International Journal of Intercultural Relations. 2004;28(5):399–414.

16. Lee J-K. Educational fever and South Korean higher education. REDIE Revista Electrónica de Investigación Educativa. 2006;8(1).

17. Bothwell E. South Korea plans ‘ghettoised’university courses for foreign students. Times Higher Education Supplement. 2015.

18. Statistical data, January 2020 [Internet]. Ministry of Justice 2020 [cited July 2020]. Available from: http://www.immigration.go.kr/immigration/1569/subview.do.

19. Moon M, Kim K-S. A case of Korean higher education reform: The Brain Korea 21 Project. Asia Pacific Education Review. 2001;2(2):96–105.

20. Yoon Y. S. Korea’s R&D Spending to GDP Ratio Highest in the World. BusinessKorea. 28 Nov 2018 [cited July 2020]. Available from: http://www.businesskorea.co.kr/news/articleView.html?idxno=26955

21. Jon JE. ‘Interculturality’ in higher education as student intercultural learning and development: a case study in South Korea. Intercultural Education. 2009;20(5):439–49.

22. Chang H-K, Han S-J, Yang N-Y, Yoo M-R, Ko E-J, Kim H-K, et al. Health status and resilience according to acculturation types among Chinese students in Korea. Korean Journal of Adult Nursing. 2010;22(6):653–62.

23. Jon J-E, Lee JJ, Byun K. The emergence of a regional hub: Comparing international student choices and experiences in South Korea. Higher Education. 2014;67(5):691–710.

24. Lee J, Jon J-E, Byun K. Neo-racism and neo-nationalism within East Asia: The experiences of international students in South Korea. Journal of Studies in International Education. 2017;21(2):136–55.

25. Alemu AM, Cordier J. Factors influencing international student satisfaction in Korean universities. International Journal of Educational Development. 2017;57:54–64.

26. Bhandari P. Stress and health related quality of life of Nepalese students studying in South Korea: A cross sectional study. Health and Quality of Life Outcomes. 2012;10(1):26.

27. Statistical data, August 2015. [Internet]. Ministry of Justice 2015 [cited July, 2020]. Available from: http://viewer.moj.go.kr/skin/doc.html?rs=/result/bbs/227&fn=1545290058798101

28. Sandhu DS, Asrabadi BR. Development of an acculturative stress scale for international students: Preliminary findings. Psychological reports. 1994;75(1):435–48.

29. Baker RW, Siryk B. Student adaptation to college questionnaire (SACQ). Los Angeles, CA: Western Psychological Services. 1986.

30. Abdullah MC, Adebayo AS. Relationship between demographic factors, social support and sociocultural adjustment among international post graduate students in a Malaysian public university. Journal of Educational and Social Research. 2015;5(2):87–92.

31. Alshammari H. Self-initiated expatriate adjustment in Saudi universities: The role of previous experience and marital status. International Journal of Business and Social Science. 2012;3(23).

32. Zhang J, Mandl H, Wang E. Personality, acculturation, and psychosocial adjustment of Chinese international students in Germany. Psychological Reports. 2010;107(2):511–25.

33. Hanassab S. Diversity, international students, and perceived discrimination: Implications for educators and counselors. Journal of Studies in International Education. 2006;10(2):157–72.

34. Karuppan CM, Barari M. Perceived discrimination and international students’ learning: an empirical investigation. Journal of Higher Education Policy and Management. 2010;33(1):67–83.

35. Suh HN, Flores LY, Wang KT. Perceived discrimination, ethnic identity, and mental distress among Asian international students in Korea. Journal of Cross-Cultural Psychology. 2019;50(8):991–1007.

